# The endemic plant species of Mt Kupe, Cameroon with a new Critically Endangered cloud-forest tree species, *Vepris zapfackii* (Rutaceae)

**DOI:** 10.1101/2021.06.01.446645

**Authors:** Martin Cheek, Jean Michel Onana

## Abstract

We revise and update the records of strict and near-endemic species of Mt Kupe, Cameroon respectively from 31 strict endemics in 2004, to 25 today, and with near-endemic species 30, unchanged in number but with turnover. The changes result from new collections, discoveries and taxonomic changes in the last 16 years. While 15 of the provisionally named putative endemic species have now been formally published, a further 18 have not. The majority of the 30 near-endemic species (18) are shared with the adjacent Bakossi Mts, far exceeding the numbers shared with the more distant Mt Etinde-Mt Cameroon, Rumpi Hills and Ebo forest areas (sharing three near-endemic species each with Mt Kupe). We test the hypothesis that a further one of the provisionally named putative Mt Kupe species, *Vepris* sp. 1 from submontane forest near the summit, is indeed new to science. We compare it morphologically with the two other bicarpellate high altitude Cameroon Highland tree species *Vepris montisbambutensis* Onana and *Vepris bali* Cheek, concluding that it is a new undescribed species here named as *Vepris zapfackii*. The new species is illustrated, mapped and its conservation status assessed as Critically Endangered using the 2012 IUCN standard due to habitat clearance from agricultural pressures at its sole location which is unprotected. *Vepris zapfackii* and *V. bali* appear unique in African trifoliolate species of the genus in having opposite leaves. *Vepris zapfackii* differs in having hairy petiolules and midribs and petiolules with the blade decurrent distally, narrowing towards a winged-canaliculate base (vs glabrous and petiolule long, terete), and sparsely golden hairy pistillodes and a glabrous calyx (vs densely black hairy pistillodes, and sepals hairy).

## Introduction

As part of the project to designate Important Plant Areas (IPAs) in Cameroon (also known as Tropical Important Plant Areas or TIPAs, https://www.kew.org/science/our-science/projects/tropical-important-plant-areas-cameroon), we are striving to name, assess the conservation status and include in IPAs (Darbyshire *et al.* 2017) rare and threatened plant species in the threatened natural habitat of Cameroon.

Several of these species were previously designated as new to science but not formally published in a series of checklists (see below) ranging over much of the Cross-Sanaga interval (Cheek *et al.* 2001). The Cross-Sanaga has the highest vascular plant species, and highest generic diversity per degree square in tropical Africa (Barthlott *et al.* 1996; Dagallier *et al.* 2020, respectively), including endemic genera such as *Medusandra* Brenan (Peridiscaceae, Breteler *et al.* 2015; Soltis *et al.* 2007). However, natural habitat is being steadily cleared, predominantly for agriculture. Eight hundred and fifteen species of vascular plant were Red Listed at the global level for Cameroon (Onana & Cheek 2011).

In this paper we test the hypothesis that the Mt Kupe cloud-forest tree species formerly designated as “*Vepris* sp. 1” (Cheek *et al.* 2004), is a new species to science, and we describe, characterise, and name it as *Vepris zapfackii* Cheek & Onana. The species is discussed in the context of revised and updated data on the endemic and near-endemic plant species of Mt Kupe, a known centre of plant diversity (see Discussion below).

*Vepris* Comm. ex A. Juss. (Rutaceae-Toddalieae), is a genus with 94 accepted species, 22 in Madagascar and 70 in Continental Africa with one species extending to Arabia and another endemic to India (Plants of the World Online, continuously updated). The genus was last revised for tropical Africa by Verdoorn (1926). Founded on the Flore du Cameroun account of Letouzey (1963), six new species were recently described from Cameroon (Onana & Chevillotte 2015; Cheek *et al.* 2018a; Onana *et al.* 2019), taking the total in Cameroon to 24 species, the highest number for any country globally. By comparison, neighbouring Gabon has seven species (Sosef *et al.* 2006). Many of these 24 species are endemic to western Cameroon (SouthWest and NorthWest Regions of Cameroon) and several are threatened (Onana & Cheek 2011) and in one case considered globally extinct (Cheek *et al.* 2018a), although only two currently appear on the IUCN Red List: *Vepris lecomteana* (Pierre) Cheek & T. Heller (Vulnerable, Cheek 2004) and *Vepris trifoliolata* (Eng.) Mziray (Vulnerable, World Conservation Monitoring Centre 1998). In other parts of Africa species are even more highly threatened, e.g., the Critically Endangered *Vepris laurifolia* (Hutch. & Dalz.) Lachenaud & Onana of Guinea-Ivory Coast (formerly *V. felicis* Breteler, Cheek 2017a; Lachenaud & Onana 2021).

In continental Africa, *Vepris* are easily recognised. They differ from all other Rutaceae because they have digitately (1–)3(–5)-foliolate (not pinnate) leaves, and unarmed (not spiny) stems. The genus consists of evergreen shrubs and trees, predominantly of tropical lowland evergreen forest, but with some species extending into submontane forests and some into drier forests and woodland. *Vepris* species are often indicators of good quality, relatively undisturbed evergreen forest since they are not pioneers (Cheek *et al.* 2019a). New species are steadily coming to light (Cheek *et al.* 2019a).

Species of *Vepris* in Africa extend from South Africa, e.g. *Vepris natalensis* (Sond.) Mziray, to the Guinean woodland in the fringes of the Sahara desert (*Vepris heterophylla* (Engl.) Letouzey). Mziray (1992) subsumed the genera *Araliopsis* Engl.*, Diphasia* Pierre*, Diphasiopsis* Mendonça, *Oricia* Pierre, *Teclea* Delile, and *Toddaliopsis* Engl. into *Vepris*, although several species were only formally transferred subsequently (e.g., Cheek *et al.* 2009, Onana & Chevillotte 2015). Mziray’s conclusions were largely confirmed by the molecular phylogenetic studies of Morton (2017) but sampling was limited, identifications appeared problematic (several species appear simultaneously in different parts of the phylogenetic trees) and more molecular work would be desirable. Morton studied about 14 taxa of *Vepris*, mainly those from eastern Africa. Nevertheless, characteristics of some of the former genera are useful today in grouping species. The “araliopsoid” species have subglobose, 4-locular fruit with 4 external grooves; the “oricioid” species are apocarpous in fruit; the fruits of “diphasioid” species are laterally compressed in one plane, bilocular and bilobed at the apex; while “tecleoid” species are unilocular in fruit and 1-seeded, lacking external lobes or grooves. There is limited support for these groupings in Morton’s study,

Due to the essential oils distributed in their leaves, and the alkaloids distributed in their roots, several species of *Vepris* have traditional medicinal value (Burkill 1997). Burkill details the uses, essential oils and alkaloids known from five species in west Africa: *Vepris hiernii* Gereau (as *Diphasia klaineana* Pierre), *Vepris suaveolens* (Engl.) Mziray (as *Teclea suaveolens* Engl.), *Vepris afzelii* (Engl.) Mziray (as *Teclea afzelii* Engl.), *Vepris heterophylla* (Engl.) Letouzey (as *Teclea sudanica* A. Chev.) and *Vepris verdoorniana* (Exell & Mendonça) Mziray (as *Teclea verdoorniana* Exell & Mendonça) (Burkill 1997: 651 – 653). Research into the characterisation and anti-microbial and anti-malarial applications of alkaloid and limonoid compounds in *Vepris* is active and ongoing, although sometimes published under generic names no longer in current use, e.g. Wansi *et al.* (2008). Applications include as synergists for insecticides (Langat 2011). Cheplogoi *et al.* (2008) and Imbenzi *et al.* (2014) respectively list 14 and 15 species of *Vepris* that have been studied for such compounds. A review of ethnomedicinal uses, phytochemistry, and pharmacology of the genus *Vepris* was recently published by Ombito *et al.* (2021) although the identification of several of the species listed needs checking. Most recently, Langat *et al.* (2021) have published three new acridones and reported multi-layered synergistic anti-microbial activity from *Vepris gossweileri* (I.Verd.)Mziray, recently renamed as *Vepris africana* (Hook.f ex Benth.) Lachenaud & Onana (Lachenaud & Onana 2021).

## Materials & Methods

This study is based on herbarium specimens. All specimens cited have been seen. The methodology for the surveys in which the specimens were collected is given in Cheek & Cable (1997). Herbarium citations follow Index Herbariorum (Thiers *et al.* continuously updated), nomenclature follows Turland *et al.* (2018) and binomial authorities follow IPNI (continuously updated). Material of the suspected new species was compared morphologically with material of all other African *Vepris,* principally at K, but also using material and images from BM, EA, BR, FHO, G, GC, HNG, P and YA. Specimens at WAG were viewed on the Naturalis website (https://bioportal.naturalis.nl/). The main online herbarium used during the study apart from that of WAG was that of P (https://science.mnhn.fr/all/search). Points were georeferenced using locality information from herbarium specimens. The map was made using simplemappr (Shorthouse 2010). The conservation assessment was made using the categories and criteria of IUCN (2012). Herbarium material was examined with a Leica Wild M8 dissecting binocular microscope fitted with an eyepiece graticule measuring in units of 0.025 mm at maximum magnification. The drawing was made with the same equipment using a Leica 308700 camera lucida attachment.

## Results

In the key to the *Vepris* species of Cameroon in Onana & Chevillotte (2015), *Vepris sp.* 1 (*Vepris zapfackii*) keys to *V. montisbambutensis* Onana because of its non-winged petiole, lack of pulvini on the petiolules, three, thinly coriaceous leaflets which are less than 20 × 6 cm, and which have brochidodromous venation, the lower surface marked by black points and also because of the ellipsoid fruits. Onana & Chevillotte (2015) left the question open as to whether these two taxa are conspecific or not. However, the opposite leaves, hairy stems, petiolules and midribs of *Vepris zapfackii* clearly separate it from *V. montisbambutensis*. In these respects, it closely resembles *Vepris bali* Cheek, previously unique in tropical Africa in having opposite trifoliolate leaves. However, *Vepris sp.* 1 (*Vepris zapfackii*) can be separated from that species and further from *V. montisbambutensis* using the characters in Table 1 below. These three species are alone in the high altitude *Vepris* species of the Cameroon Highlands in the bifurcate pistillodes of the male flowers (not available in *V. montisbambutensis*) and the apocarpous bicarpellate fruits (not seen in *Vepris bali*).

**Table 1.** Diagnostic characters separating montane or upper submontane *Vepris* tree species of Cameroon Highlands with bicarpellate apocarpous ovaries/pistillodes bifid and/or fruits apocarpous. Data for *Vepris montisbambutensis* from Onana & Chevillotte (2015).

|  | *Vepris montisbambutensis* | *Vepris bali* | *Vepris zapfackii* |
| --- | --- | --- | --- |
| Phyllotaxy | Leaves alternate | Leaves opposite | Leaves opposite |
| Petiolule shape | Terete (cylindric) | Terete (cylindric) | Winged-canaliculate |
| Petiolule length (mm) | 2 – 10 | 9 – 14 | (1 – )2 – 6( – 7) |
| Indumentum of stems | Glabrous | Patent puberulent | Appressed hairy |
| Indumentum of petioles | Glabrous | Glabrous | Simple, spreading, brown-hairy |
| Petiole shape | Terete (cylindric) | Terete (cylindric) | Canaliculate |
| Leaflet apex | Shortly acuminate, acumen to 0.5 cm, entire | Shortly acuminate, acumen 0.1 – 0.35 cm, entire (rarely bifid), lacking a | Shortly acuminate acumen retuse, (0.2 – )0.3 – 0.5( – 0.6) cm with a central |
|  |  | central protrusion of the midrib | protrusion of the midrib. |
| Pistillode of male flower: indumentum | Not known | Surface completely covered in black, bristle like hairs | Surface partly covered (c. 50% of surface) in scattered golden hairs. |
| Sepal margin indumentum | Not known | Densely pectinate-hairy | Glabrous |
| Median leaflet dimensions (cm) | (2 – )3 – 9 x (0.7 – )1.5 – 3 | 6 – 10.5 x 2.8 – 4.7 | (4.5 – )6 – 9.5( – 10) x (1.8– )2.8 – 3.6( – 4.4) |

### Vepris zapfackii

*Cheek & Onana* **sp. nov.** Type: Cameroon, South West Region, Mt Kupe, above Nyasoso, fr. June 1996, *Zapfack* 1005 (holotype K001394679; isotypes B, ETH, US, YA) (Fig. 1).

**Fig. 1.**
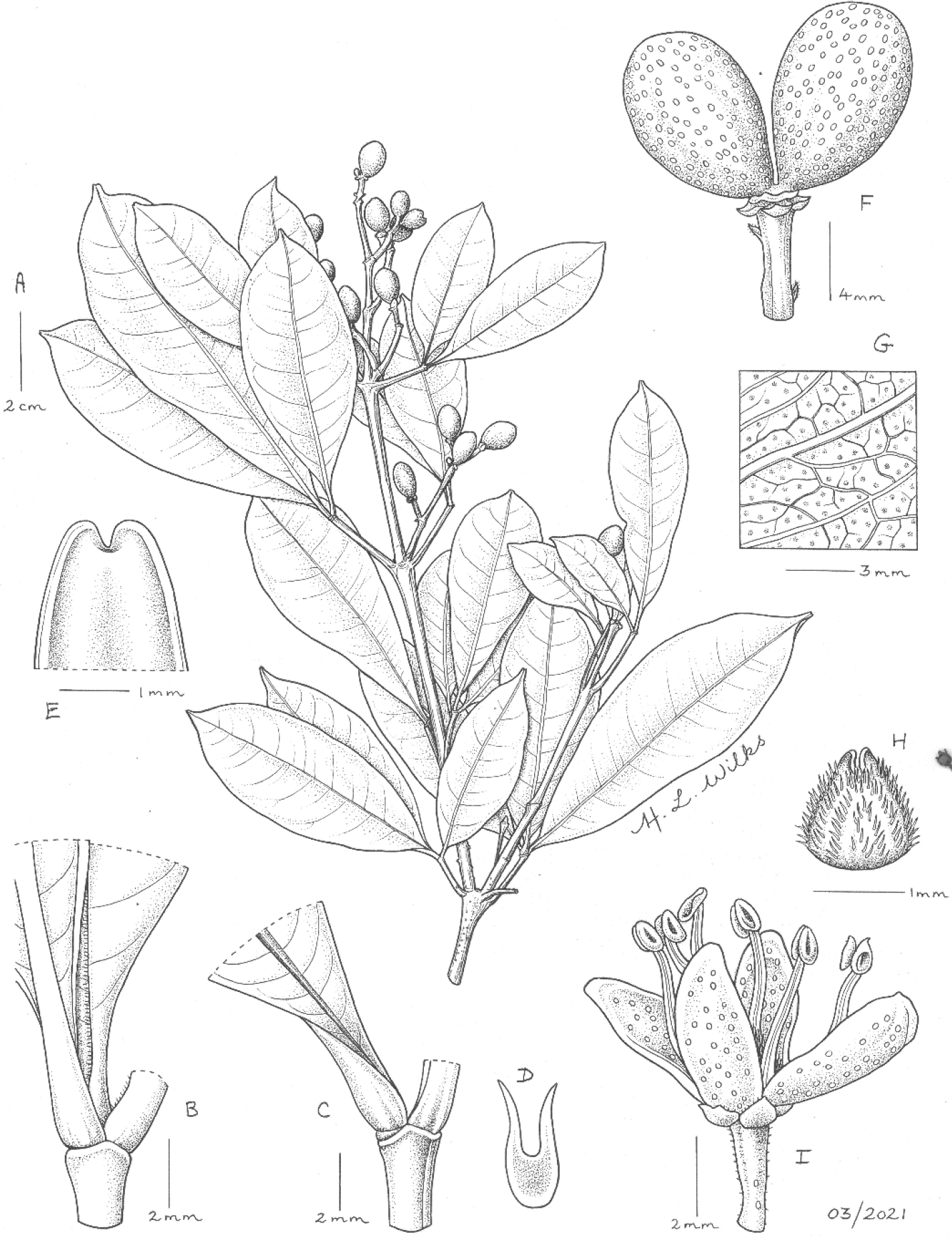
*Vepris zapfackii.* **A.** habit, fruiting branch; **B.** indumentum of leaflets; **C.** base of leaflets showing decurrent-winged canaliculate petiolules; **D.** Transverse section of petiolule-proximal part; **E.** retuse acumen; **F.** fruit with two apocarpous mericarps; **G.** gland dots on leaf abaxial surface; **H.** pistillode (male flower) side view; **I.** male flower, side view. **A-G** from *Zapfack* 1005 (K); **H** & **I** from *Ryan* 407, drawn by HAZEL WILKS

Syn. *Vepris sp.* 1 Cheek *et al.* (2004: 396).

*Monoecious or dioecious* evergreen shrub or tree 1 – 6 m tall. Trunk characteristics not recorded. Leafy stems terete, 3 mm diam. at most proximal leafy node, internodes (1 −)1.5 – 3.6(− 5.5) cm long, epidermis drying yellow-green, longitudinally fluted, moderately densely minutely appressed hairy, hairs simple, golden-brown, 0.05(– 0.1) mm long, at stem apex, covering c. 60% of the surface, gradually decreasing in density with distance from apex, c. 50% cover at third node from apex. *Leaf* phyllotaxy opposite and decussate, leaves with pleasant scent when crushed (*Sebsebe* 5097) trifoliolate, coriaceous, drying glossy green, 8.5–12 cm long, median leaflet oblanceolate, lateral leaflets 75 – 85% the length of the median, narrowly elliptic, elliptic-rhombic, or elliptic-oblanceolate, (4.5 −)6 – 9.5(− 10) × (1.8 –)2.8–3.6(– 4.4) cm, apex shortly but distinctly acuminate, acumen (0.2 −)0.3 – 0.5(– 0.6) cm long, apex of acumen minutely retuse, the cleft 0.5 mm deep, the cleft edges parallel, curving from the horizontal plane by 90 degrees abaxially, forming a V-shape, the vertex abaxial, the margins thickened, revolute. (Fig. 1E), base of leaflet acute, decurrent as a wing to the distal half of the petiolule (Fig. 1C); margin thickened, revolute; secondary nerves, with midrib and tertiary nerves, raised, thickened, (5 −)6 – 8 on each side of the midrib, yellow-green, arising at 90° from the midrib, arching towards the leaflet apex, forming a looping infra-marginal nerve c. 2 mm from the margin with a parallel, more minor infra-marginal nerve <1 mm from the margin. Intersecondary nerves 1(– 3), patent, extending towards margin before branching. Tertiary nerves inconspicuous, sparse; essential oil glands (5 −)6(− 7) per mm^2^, orbicular, 0.2 mm diam., appearing black on the abaxial surface (Fig. 1G); surfaces glabrous, except midrib (Fig. 1B) and near the petiolule, indumentum as petiolule (below); petiolules winged broadly distally, the wings becoming progressively narrower and erect in the proximal part, and laterally compressed and deeply canaliculate (Fig. 1D), (0.1 −)0.2 – 0.6(− 0.7) cm long, with scattered simple, brown hairs 0.05 mm long (Fig. 1B), glabrescent. Petiole articulated with the petiolules of the three leaflets, shallowly canaliculate, (1.2 −)1.4 – 2(− 2.9) cm long, 1.05 – 1.4 mm wide, hairs scattered, brown, 0.05 mm long, spreading, glabrescent. *Male inflorescences* (Female unknown) terminal, or in axillary pairs of the most distal nodes of the leafy stems, 5.5 – 7 × 3 – 4.5 cm, 40 – 50-flowered, peduncle 1.5 – 2.4 cm long; partial-peduncles opposite, in 2 – 4 pairs along main axis, 1.2 – 1.7 cm long, rhachis internodes 3 – 5, 3 – 10 mm long; partial-inflorescences 1.5 – 3.5 cm long, bracts patent, narrowly triangular 0.5 – 0.55 mm long, margin with dense, long, translucent hairs, bracteoles 2, subtended by the bracts, similar but smaller, c. 0.45 mm long. Pedicels (1 −)1.5 2(− 2.8) mm long, scattered with patent hairs 0.05 mm long. *Flowers* white, c. 3 mm wide (Fig. 1I). Sepals 4, broadly ovate or quadrate, slightly spreading and concave, c. 0.5 × 0.75 mm, apex acute to obtuse, glabrous. Petals 4, elliptic-oblong, 4.5 – 4.9 × 1.8 – 2 mm, apex rounded, lacking appendages; oil glands orange, elliptic or orbicular, 0.1 – 0.2 mm diam., 7 – 20 scattered on abaxial and adaxial surfaces except for the scarious, 0.2 mm wide, very minutely hairy margin. Stamens 8, equal, slightly exceeding petals, filaments 2.5 – 4(– 4.5) mm long, ribbon-like; anthers ellipsoid, 0.75(– 1) × 0.5 mm. Pistillode (vestigial ovary), broadly ovoid to conical 1.25 × 1.25 mm, apex bifurcate, cleft 0.25 mm deep, drying black, with sparse to moderately dense spreading golden hairs 0.1 – 0.15 mm long covering c. 50% of the surface (Fig. 1H). *Fruit* (immature) apocarpous, mericarps 2 (sometimes 1 by abortion), barely united at the base (Fig. 1F), obovoid to ellipsoid, 0.9 – 1.0 × 0.5 – 0.7 cm, glabrous, surface with oil glands densely scattered, elliptic, 0.75 × 0.5 mm, apex rounded, apiculus absent.

### RECOGNITION

*Vepris zapfackii* shares many similarities with *Vepris bali* Cheek, the two sharing the very unusual character in tropical African *Vepris* of opposite trifoliolate leaves. Both species also share a terminal paniculate inflorescence, and a hairy, bifid pistillode in the male flowers. They differ in that the petiolule of *Vepris zapfackii* is winged to deeply canaliculate throughout its length (vs terete), the petiole canaliculate (vs terete), the petioles, petiolules and midribs are sparsely hairy (vs glabrous), the sepals are glabrous (vs hairy), the pistillode is incompletely (c. 50% of the surface) covered in golden bristle hairs (vs completely covered in black hairs). In addition to the differential characters given in the diagnosis and Table 1, *Vepris zapfackii* also differs from *Vepris bali* in that the lateral leaflets are 75 – 80% the length of the median leaflet (vs c. 50%), the median leaflets are usually narrowly elliptic or oblanceolate (not obovate), and the leaflet apices are usually minutely retuse, the sides of the cleft oriented at 90 degrees from the plane of the remainder of the leaflet (vs rarely retuse) and the midrib projects into the cleft as a small peg (not illustrated). It additionally differs from *V. montisbambutensis* in that the surface of the fruits is not foveolate but smooth, the gland dots are minute. However, it seems very likely that the three are sister species. *Vepris montisbambutensis* and *V. bali* occur to the north, and are separated by about 100 and 130 km, respectively, from *Vepris zapfackii* along the length of the Cameroon Highlands. *Vepris montisbambutensis* is a point endemic restricted to the area of the Lebialem Highlands/Bamboutos Mts, and *Vepris bali* is, or was, endemic to high altitude in the Bali Ngemba Forest Reserve. Additional differential characters are given in Table 1

### DISTRIBUTION

Cameroon, known only from the summit of Mt Kupe (at the border of South West and Littoral Regions) (Map 1).

### SPECIMENS EXAMINED. CAMEROON

**South West Region, Mt Kupe**, above Nyasoso, fr. June 1996, *Zapfack* 1005 (B, ETH, K001394679, US, YA); ibid., above Kupe Village, 1940 m alt., Top of Kupe Mt, male fl. 30 May 1996, *Ryan* 407 (K000198298, MO, P, SCA, WAG, YA); ibid., towards the summit (peak 1) eastwards from the base camp (No. 2), immature fr., 11 Feb 1995, *Sebsebe* 5097 (K000198299, MO, P, WAG, YA).

### HABITAT

Upper submontane forest dominated by *Garcinia* with many Rubiaceae (*Sebsebe* 5097), also with *Zenkerella citrina* Taub. c. 1940 m alt.

**Map 1.**
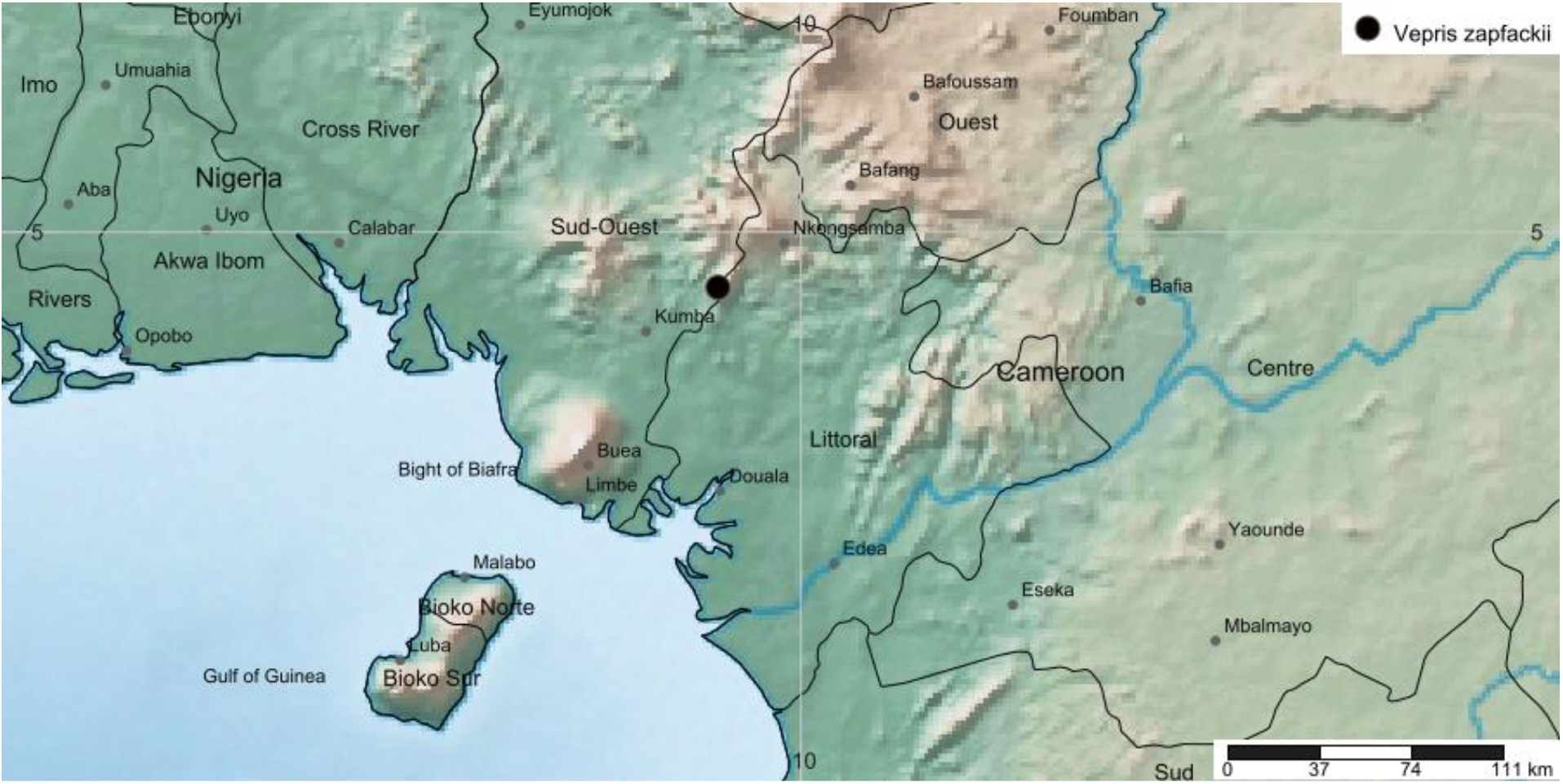
Global distribution of *Vepris zapfackii.*

### CONSERVATION STATUS

*Vepris zapfackii* is known from only three specimens all of which are confined to the forested summit area of Mt Kupe (2064 m), from personal experience at this location, the first author estimates the number of mature individuals of the species to be <50.

Mt. Kupe is not a formally protected area. The summit area is estimated to be <4 km^2^. Clearance of forest for agriculture is continuing upslope at Mt Kupe. The greatest advance has occurred on the east side where it is already close to the summit. On the west side clearance has not yet been observed above 1500 m altitude but since there are no restrictions in place, the threat to rare, range-restricted species dependent on intact forest for survival is evident. Accordingly, we here assess *Vepris zapfackii* as Critically Endangered. CR B1+B2ab(iii)+D.

It is possible that *Vepris zapfackii* will yet be found at additional locations in Cameroon. However, while surveys have not been exhaustive, many thousands of specimens have been collected in areas to the north, south, west and east of Mt Kupe, yet no additional specimens of this distinctive species have been found (Cheek *et al.* 1996; Cable & Cheek 1998; Cheek *et al.* 2000; Maisels *et al.* 2000, Chapman & Chapman 2001; Harvey *et al.* 2004; Cheek *et al.* 2004; Cheek *et al.* 2006; Cheek *et al.* 2010; Harvey *et al.* 2010; Cheek *et al.* 2011).

### PHENOLOGY

Flowering in late May, immature fruits in February and June.

### ETYMOLOGY

Named for Dr Louis Zapfack (1960 –) who collected the type specimen of this species. Raised in Dschang, West Region, he is currently Professor in the Department of Plant Biology, Faculty of Science, University of Yaoundé I, where he is an ecologist and Climate Change Specialist (Carbon Stock evaluation). In the mid-1990s he led volunteer teams under the Earthwatch programme, linked with the “Plant Diversity of Western Cameroon” programme (Darwin Initiative), collecting many hundreds of herbarium specimens, including *Vepris zapfackii*. He is also commemorated by *Bafutia tenuicaulis* var. *zapfackiana* Beentje & B.J. Pollard in Cheek *et al.* (2000: 92).

### VERNACULAR NAMES & USES

None are known.

## Discusssion

### The Cameroon Highlands and Mt Kupe

The Cameroon Highlands extend through four tropical African countries, along a fault line between two major African plates. Beginning in the south on the volcano island of Bioko (Equatorial Guinea) they continue on the mainland with the isolated Mount Cameroon active volcano (at 4095 m the highest point of the highlands), then heading 110 km NNE forming the ridges, plateaux and isolated peaks of the Bakossi Mts and Mts Kupe, Muanenguba, continuing NNE to the Ntale Mts of Banyang Mbo, the Lebialem Highlands and Bamboutos Mts, the Bamenda Highlands including Mt Oku (3011 m), Tchabal Mbabo, then heading eastwards and forming the lower and drier Adamaoua Highlands which extend into the Central African Republic. Two westward extending arms from the central section in Cameroon extend into Nigeria, forming the Obudu and Mambilla Plateaux. Four main mountain building periods, three volcanic and one plutonic have been reviewed by Courade (1974). Mt Kupe (2064 m) is mainly formed of syenite and granite with a basaltic cap (Enang *et al.* 2020) uplifted in the third, plutonic period, as was probably Mt Nlonako (1790 m) c. 30 km to the NE. The more extensive Bakossi Mts (1895 m) to the west, are uplifted pre-Cambrian basement complex punctured by two small volcaniccraters, and are separated from Mt Kupe by the 5 – 10 km wide Chide trough fault. To the west of the Bakossi Mts, separated by the valley of the Mungo River, the Rumpi Hills are formed from ancient basalts and trachytes. The Chide trough, with its volcanic cinder cones producing fertile soils for agriculture, is comparatively densely populated. 25 km to the north of Mt Kupe is the 3 – 4 km wide Mwanenguba caldera (2411 m) occupying the summit of a massive Hawaiian type volcano.

Mt Kupe is surrounded by a 4 m p.a. isohyet in Courade (1974) but rainfall varies from 3 m p.a. at the town of Loum on its SE corner, to 6 – 7 m p.a on the slopes of the SW (Sieffermann 1973 cited in Cheek *et al.* 2004). The main rainy season is May-October inclusive. All but two months, December and January have rainfall exceeding 100 mm at Tombel in the SW and Nyasoso in the W where relative humidity never falls below 80%. However, measurements have only been made at settlements below and not on the mountain itself. Throughout the year the mountain is blanketed in cloud and there are only rare moments when this is lifted, such as immediately after a heavy thunderstorm. Submontane, also known as cloud forest is generally accepted to occupy the 800 – 2000 m altitudinal interval and covers the mountain in that range apart from some elements of montane forest and a few grassy spots at the summit where soil is too thin over rock surfaces to support trees, and also several large, vertical cliff faces, locally known as “rocks”, which have a distinct community of plant species. Lowland forest at the 400 – 600 m foot of the mountain was mainly evergreen, but semi-deciduous in the SE. It has mainly been cleared for agriculture, especially coffee, *Coffea canephora*, Pierre ex Froehn., cacao, *Theobroma cacao* L., cabbage, *Brassica oleracea* L. and plantain *Musa* spp. (Stoffelen *et al.* 1997). However, remnants occur, with several point plant species endemics.

Twelve Bakossi villages surround the c. 42 km2 forest of Mt Kupe. Traditionally the summit of the mountain is considered the home of powerful spirits that can enslave humans, and is regarded as a fearsome and sacred place, and it is well-known throughout Cameroon for this reason (Wild in Cheek *et al.* 2004). However, a trail to the summit from Nyasoso (800 m) on the west side has been popular with ecotourists, especially those seeking to view the submontane bird species, the Mt Kupe Bush Shrike (*Chlorophoneus kupeensis,* formerly placed in *Malaconotus* and *Telophorus*) previously considered endemic to Mt Kupe but now known to occur also in the Bakossi Mts to the west and at two points in adjoining Nigeria. Less than 250 individuals are estimated to exist and it is assessed as Endangered (Birdlife International 2018).

Mt Kupe and the Bakossi Mts, together with the slopes of Mwanenguba, and the adjoining Ntale probably comprise the largest surviving area of submontane forest in both Cameroon and in Africa west of the East African rift mountains. Elsewhere in the Cameroon Highlands this vegetation has either been completely cleared for agriculture (the Bamileke Plateau and Bamboutos Mts) or reduced to small fragments (Lebialem, Bamenda and Mambilla Highlands) or due to lower humidity at higher latitudes, it appears absent as at Tchabal Mbabo and the Adamaoua Mts (Chapman & Chapman 2001; Harvey *et al.* 2004; Harvey et al 2010). However, cloud forest also occurs on Mt Cameroon (Tchouto *et al.* 1999).

### The endemic plant species of Mt Kupe

Unlike the well-known Mt Cameroon (Cheek *et al.* 1994; Cheek *et al.* 1996), Mt Kupe and the neighbouring Bakossi Mts were unknown to be centres of plant species diversity until the early 21^st^ Century when the “Conservation Checklist for the Plants of Kupe, Mwanenguba and the Bakossi Mountains” was published (Cheek *et al.* 2004). In one of the first papers foreshadowing this and reviewing knowledge of the rare species then known from Mt Kupe while describing a species then considered endemic to the mountain (*Coffea montekupensis* Stoff., Stoffelen *et al.* 1997), no other endemic species were noted although five rare and near-endemic species were listed and six new species were stated as being in the course of preparation for publication (Stoffelen *et al.* 1997). After the collection of many thousands of herbarium specimens in the late 1990s (including the specimens described in this paper), and subsequent identifications, large numbers of new species such as *Vepris* sp. 1, came to light and were recorded in Cheek *et al.* (2004).

Twenty-six strict endemics to Mt Kupe and 30 near endemics were recorded there from submontane forest (Cheek *et al.* 2004: 37 – 39) while three further strict endemics were recorded from montane habitat (Cheek *et al.* 2004: 58 & 65) and two from lowland habitats (Cheek *et al.* 2004: 29). Many of the endemics were known under provisional names only. Since that time, many of these have been formally published. Some of the species previously thought to be endemic have been discovered at additional locations so that they are no longer either endemic to Mt Kupe (known only from Mt Kupe) or near-endemic (here defined as from Mt Kupe and one or two other locations). Examples of species previously considered endemic but no longer so are: *Coffea montekupensis* Stoffeln. (Stoffelen *et al.* 1997) now recorded not only from Mt Kupe and the Bakossi Mts, but from Ebo, Banyang Mbo and the Lebialem Highlands (Harvey *et al.* 2010:135). Similarly, *Uvariopsis submontana* Kenfack, Gosline & Gereau (Kenfack *et al.* 2003) is now known from more than two locations apart from Mt Kupe although still rare and threatened. *Mussaenda* sp. nov. later published as *M. epiphytica* Cheek (Cheek 2009) is now also recorded from Mt. Etinde, Rumpi Hills & Mbam Minkom (Lachenaud *et al.* 2013). *Tricalysia* sp. B *aff. ferorum,* published as *Tricalysia elmar* Cheek is now also recorded from Ngovayang & Rumpi Hills (Cheek *et al.* 2020a). *Psychotria sp. aff. camerunensis* of Cheek *et al.* (2004), published as *P. ngollengollei* Cheek (Cheek *et al.* 2009) was reduced to a subspecies of *P. solfiana* K. Krause and shown also to be at Rumpi Hills, Banyang Mbo and Nlonako, apart from Mt. Kupe and Bakossi Mts (Lachenaud 2019). *Dracaena* sp. aff. *phrynoides* appears to be *Dracaena bushii* Damen with nine locations (Damen *et al.* 2018).

Other taxa formerly thought to be possibly new to science have subsequently been reidentified e.g. *Friesodielsia enghiana* var. nov. as *Monanthotaxis glaucifolia* (Hutch. & Dalz.) P.H. Hoekstra, *Monanthotaxis* sp. nov. as *Sphaerocoryne gracilipes (*Beath.) X. Guo & R.M.K. Saunders (Couvreur *et al.* 2021), and *Bulbophyllum kupense* P.J. Cribb & B.J. Pollard (Cribb & Pollard 2004) has been reduced to synonymy of *B. teretifolium* Schltr., still rare and threatened but recorded from Mbam Minkom, Rumpi Hills and Banyang Mbo in addition to Mt Kupe, so is no longer a near endemic (Droissart *et al.* 2012).

Below, in Table 2, we update the known endemics of Mt Kupe. In addition to those species originally documented as such in Cheek *et al.* (2004) often now under other names, we include species that had not been included there, but which have come to light subsequently, such as *Psychotria asterogramma* O. Lachenaud (Lachenaud 2019:150) endemic to Mt Kupe, Mt Cameroon and Bioko and *Aframomum kamerunicum* D.J. Harris & Wortley, endemic to Mt Kupe and the Bakossi Mts (Harris & Wortley 2018).

**Table 2.** The strictly endemic plant species of Mt. Kupe, and those near-endemic, i.e. also at one or two other locations: the Bakossi Mts, Mt Nlonako, Lebialem Highlands or Ebo. Revised and updated from Cheek *et al.* (2004: 37 – 39).

| Current name | Previously known as (e.g. in Cheek *et al.* (2004)) | Strictly Endemic to Mt Kupe | Also Bakossi Mts | Also Rumpi Hills | Also Nlonako | Also Mt Cameroon-Etinde | Also Lebiale Highlands | Also Ebo | Reference |
| --- | --- | --- | --- | --- | --- | --- | --- | --- | --- |
| *Diospyros kupensis* |  |  | ✓ |  |  |  |  |  | Gosline & Cheek (1998) |
| *Drypetes* sp.A |  |  | ✓ |  |  |  |  |  |  |
| *Eugenia* sp. 1 |  |  | ✓ |  |  |  |  |  |  |
| *Coffea bakossii* |  |  | ✓ |  |  |  |  |  | Cheek *et al.* (2002a) |
| *Ixora* sp. A |  |  | ✓ |  |  |  |  |  |  |
| *Lasianthus* sp. A |  |  | ✓ |  |  |  |  |  |  |
| *Pentaloncha* sp. nov. |  |  | ✓ |  |  |  | ✓ |  |  |
| *Psychotria bakossiensis* | *Psychotria* sp. A aff. *gabonica* |  | ✓ |  |  |  |  |  | Cheek & Sonké (2005a) |
| *Sericanthe* sp. A |  |  | ✓ |  |  |  |  |  |  |
| *Tricalysia* sp. F |  |  | ✓ |  |  |  |  |  |  |
| *Rhaptopetalum geophylax* |  |  | ✓ |  |  |  | ✓ |  | Cheek *et al.* (2002b) |
| *Cola etugei* | *Cola* sp. nov. 'etugei' | ✓ |  |  |  |  |  |  | Cheek *et al.* (2020b) |
| *Dracaena kupensis* | *Dracaena* cf. *phanerophlebia* |  | ✓ |  |  | ✓ |  |  | Mwachala *et al.* (2007) |
| *Brachystephonus kupeensis* |  | ✓ |  |  |  |  |  |  | Champluvier & Darbyshire (2009) |
| *Justicia* sp. 1 (Ngomboku) |  | ✓ |  |  |  |  |  |  |  |
| *Drypetes* sp. B |  | ✓ |  |  |  |  |  |  |  |
| *Memecylon kupeanum* | *Memecylon* sp. 1 | ✓ |  |  |  |  |  |  | Stone *et al.* (2008) |
| *Ardisia koupensis* |  |  |  |  | ✓ |  |  |  |  |
| *Ardisia alabastroalatum* |  | ✓ |  |  |  |  |  |  |  |
| *Eugenia* sp. 2 |  | ✓ |  |  |  |  |  |  |  |
| *Kupeantha kupensis* | *Calyco-siphonia* sp. A | ✓ |  |  |  |  |  |  | Cheek *et al.* (2018b) |
| *Ixora* sp. B |  | ✓ |  |  |  |  |  |  |  |
| *Psychotria* sp. aff. *dorotheae* |  | ✓ |  |  |  |  |  |  |  |
| *Psychotria* sp. B aff. *leptophylla* |  | ✓ |  |  |  |  |  |  |  |
| *Rytigynia* sp. B |  | ✓ |  |  |  |  |  |  |  |
| *Rytigynia* sp. C |  | ✓ |  |  |  |  |  |  |  |
| *Tricalysia* sp. C |  | ✓ |  |  |  |  |  |  |  |
| *Vepris zapfackii* | *Vepris* sp. 1 | ✓ |  |  |  |  |  |  | (this paper) |
| *Pancovia* sp. 1 |  | ✓ |  |  |  |  |  |  |  |
| *Microcos magnifica* | *Microcos* sp. A |  |  |  |  |  |  | ✓ | Cheek (2017b) |
| *Afrothismia kupensis* | *Afrothismia pachyantha* s.l. | ✓ |  |  |  |  |  |  | Cheek *et al.* (2019b) |
| *Afrothismia saingei* |  | ✓ |  |  |  |  |  |  | Franke (2004) |
| *Afrothismia fungiformis* |  |  |  |  |  |  |  | ✓ | Sainge *et al.* (2013) |
| *Costus kupeensis* | *Costus* sp. A |  |  |  |  |  |  | ✓ | Maas-van der Kamer *et al.* (2016) |
| *Cyperus tenuiculmis* subsp. *mutica* |  | ✓ |  |  |  |  |  |  |  |
| *Bulbophyllum jaapii* |  | ✓ |  |  |  |  |  |  | Szlachetho & Olsewski (2001) |
| *Polystachya kupensis* |  | ✓ |  |  |  |  |  |  | Cribb & Pollard (2002) |
| *Kupea martinetugei* |  |  |  |  |  | ✓ |  |  | Cheek *et al.* (2003) |
| *Aframomum kamerunicum* |  |  | ✓ | ✓ |  |  |  |  | Harris & Wortley (2018) |
| *Cyperus microcristatus* |  | ✓ |  |  |  |  |  |  | Lye & Pollard (2006) |
| *Psydrax bridsoniana* |  | ✓ |  |  |  |  |  |  | Cheek & Sonké (2005b) |
| *Psychotria darwiniana* | *Psychotria* sp. aff. *alatipes* |  | ✓ | ✓ |  |  |  |  | Cheek *et al.* (2009) |
| *Pavetta kupensis* |  |  | ✓ |  |  |  |  |  | Manning (1996) |
| *Lefebvrea kupense* | *Peucedanum kupense* | ✓ |  |  |  |  |  |  | Cheek *et al.* (2004) |
| *Dissotis* sp. 1 |  | ✓ |  |  |  |  |  |  |  |
| *Psychotria cheekii* | *Psychotria* sp. 9 |  |  | ✓ |  |  |  |  | Lachenaud (2019) |
| *Psychotria asterogramma* |  |  |  |  |  | ✓ |  |  | Lachenaud (2019) ((also Bioko) |
| TOTALS |  | 25 | 15 | 3 | 1 | 3 | 2 | 3 |  |

Examination of Table 2 shows that the number of strict endemics on Mt Kupe is now 25, a drop of six from the 31 in Cheek *et al.* (2004). This change has resulted from further survey work in other locations which has extended the known ranges of the species concerned so that they are no longer strict endemics. It has also resulted from taxonomic revisions and floristic studies that have reduced to synonymy three of the previously supposed strict endemics.

The number of near-endemics for Mt Kupe at 30, remains the same as in 2004. However, there has been some turnover with replacement on the list of formerly supposed near endemics since found to have wider ranges, with both former strict endemics now found to be near endemics and four newly published near endemic taxa that had not been listed as endemics in 2004, even under provisional names.

By far the majority of near-endemics (18/30) of Mt Kupe are shared with the Bakossi Mts only 5 – 10 km to the west. This number can be expected to increase if botanical surveys are continued since the Bakossi Mts are far from comprehensively studied: much of its vast area remains unvisited by botanists. Yet the Bakossi Mts have many endemics that do not extend to Mt Kupe, e.g. *Impatiens frithii* Cheek (Cheek & Csiba 2002). Those near-endemics shared with the more distant Mt Cameroon and Rumpi Hills, only three each, reflect their greater geographical separation. Unexpectedly an equal number of near-endemics (3) are shared with the Ebo Forest, not formerly considered part of the Cameroon Highlands, c. 100 km to the SSE and which also has its own strict endemics, documented in Cheek *et al.* 2018c, with new additions currently being added (Gosline *et al.* 2021; Cheek *et al.* 2021). The very unexpected low number (1) shared with Mt Nlonako likely reflects low survey effort at this location.

While 15 of the provisionally named supposed endemics of Cheek *et al.* (2004) have now been formerly published (11 of which remain endemic or near-endemic) a further 18 (of which 11 are strictly endemic) remain unpublished. This paper continues the effort to address this deficit.

Formal publication of provisionally named endemics is a high priority since until species receive a scientific name, it is difficult for their conservation assessments to be placed on the IUCN Red List. Although formal protection of Mt Kupe was thought to be imminent in 2004 (Cheek *et al.* 2004), this has proved not to be the case. Instead, forest clearance for agriculture has continued increasing upslope reaching 1500 m alt. in some places (Birdlife International 2018). As such, the endemic plant species that depend upon it must all be considered threatened.

## Conclusions

About 2000 new species of vascular plant have been discovered by science each year for the last decade or more adding to the estimated 369 000 already known (Nic Lughadha *et al.* 2016), although the number of flowering plant species known to science is disputed (Nic Lughadha *et al.* 2017). Until species are delimited and known to science, it is more difficult to assess them for their conservation status and so the possibility of protecting them is reduced (Cheek *et al.* 2020c). In the main, the majority of new species described today tend to be range-restricted, making them especially likely to be threatened, although there are some exceptions (e.g. Cheek & Etuge 2009). To maximise the survival prospects of range-restricted species there is an urgent need not only to document them formally in compliance with the requirements of the relevant nomenclatural code (Turland *et al.* 2018), but also to formally assess the species for their extinction risk, applying the criteria of a recognised system, of which the IUCN Red List of Threatened Species is the most widely accepted (Bachman *et al.* 2019). Despite rapid increases over recent years in numbers of plant species represented by assessments on the Red List, the vast majority of plant species still lack such assessments (Nic Lughadha *et al.* 2020). Documented extinctions of plant species are increasing (Humphreys *et al.* 2019) and recent estimates suggest that as many as two fifths of the world’s plant species are now threatened with extinction (Nic Lughadha *et al.* 2020). In Cameroon *Oxygyne triandra* Schltr. is globally extinct as is *Afrothismia pachyantha* Schltr. (Cheek & Williams 1999; Cheek *et al.* 2018d; Cheek *et al.* 2019b), and at least two species of the African genus *Inversodicraea* (Cheek *et al.* 2017). In some cases, species appear to be extinct even before they are named for science, such as *Vepris bali* Cheek (Cheek *et al.* 2018a), and in neighbouring Gabon, *Pseudohydrosme bogneri* Cheek & Moxon-Holt ined. (Moxon-Holt & Cheek 2021). Most of the >800 Cameroonian species in the Red Data Book for the plants of Cameroon are threatened with extinction due to habitat clearance or degradation, especially of forest for small-holder and plantation agriculture e.g. oil palm, following logging (Onana & Cheek 2011). Efforts are now being made to delimit the highest priority areas in Cameroon for plant conservation as Tropical Important Plant Areas (TIPAs) using the revised IPA criteria set out in Darbyshire *et al.* (2017).

National governments and leaders have recognised the importance of species assessed as threatened on the Red List and documented in IPAs or TIPAs as demonstrated recently in Cameroon when in part due to the high number of plant species on the Red List for the Ebo TIPA (Lovell 2020), a logging concession was revoked for the Ebo forest (Kew Science News 2020).

It is hoped that formal publication and Red Listing of additional endemic species such as the Critically Endangered *Vepris zapfackii* will help support the case for Mt Kupe being evidenced as a TIPA and gaining formal protection to help avoid the risk of global extinction of the 55 endemic plant species currently documented there, of which 25 are globally unique to the mountain on current evidence.

## Acknowledgements

The specimens cited in this paper were collected by or with the support of volunteers and sponsored scientists arranged by Earthwatch Europe, Oxford, including Sebsebe Demissew, Louis Zapfack, P. Ryan, C. Moloney, and G. Adamu, and we were also assisted greatly by our Bakossi colleagues such as Daniel Ajang and the late Martin Etuge. George Gosline and Stuart Cable of Royal Botanic Gardens, Kew gave invaluable support at Mt Kupe as did Chris and Liz Bowden of the former Mt Kupe Forest Project (Birdlife International). Drs Satabie, Achoundong and more recently Florence Ngo Ngwe, Eric Nana, Jean Betti Lagarde, the current and former directors, of IRAD-National Herbarium of Cameroon, Yaoundé, and their staff are thanked for expediting the collaboration between our two institutes. Janis Shillito typed the manuscript. Two anonymous reviewers are thanked for constructively reviewing an earlier version of this paper.

